# Comparative spatial transcriptomics of hair follicle-T cell interactions in mouse, dog and human reveals conserved drivers of primary cicatricial alopecia

**DOI:** 10.1101/2025.01.20.633953

**Authors:** Ümmügülsüm Yıldız-Altay, Rutha Adhanom, Warda Abdi, Andrew Romasco, Lauren You, Elise Santacruz, Saeed Shakiba, Mridushi Daga, Qi Tang, Matthew Olivera, Thomas Cicuto, Floriane Bretheau, Christina Baer, Motahareh Arjomandnejad, Allison Keeler, Helen Bui, Babak Shokrani, Ramon Almela, Angel S. Byrd, Cheri Frey, Jillian M. Richmond

**Affiliations:** Department of Dermatology, UMass Chan Medical School, Worcester, MA, USA; RNA Therapeutics Institute, UMass Chan Medical School, Worcester, MA, USA; SCOPE, UMass Chan Medical School, Worcester, MA, USA; Bliq Photonics, Quebec City, Quebec, Canada; Gene Therapy Center, UMass Chan Medical School, Worcester, MA, USA; Department of Dermatology, Howard University College of Medicine, Washington, DC, USA; Department of Pathology, Howard University College of Medicine, Washington, DC, USA; Department of Clinical Sciences, Tufts Cummings School of Veterinary Medicine, North Grafton, MA, USA

**Keywords:** discoid lupus erythematosus (DLE), primary cicatricial alopecia (PCA), lymphocytic inflammation, hair follicle, immune privilege collapse, CD8+ T cell, digital spatial profiling (DSP), spatial transcriptomics

## Abstract

Primary cicatricial alopecias (PCA) encompass several autoimmune disorders characterized by scarring hair loss. Many of these conditions are lymphocytic and are thought to be driven by T cell populations. Here, we sought to characterize potential T cell-hair follicle communication pathways in the microanatomical niche using spatial transcriptomics across 3 mammalian species including a novel mouse model, spontaneous disease in companion dogs and human archival diagnostic biopsies. Flow cytometry of mouse model skin confirmed loss of CD34+ bulge cells and keratinocytes, and bulk microarray and histology revealed expression of collagens and development of fibrosis. *In vivo* ear imaging in mice engrafted with Kikume photoconvertible OT1 CD8+ T cells confirmed long-lived RFP+ T cells in skin arrest near hair follicles and recruit other GFP+ T cells. OT1 T cells expressed CD69, CD103, CD122 and CD62L, which is a binding partner of CD34. Digital spatial profiling using CD3, CD8 and CD45 cell masking identified CXCR3 ligands and IFN response genes in hair follicles, and “metabolic” pathways in T cells, which were also recapitulated in dog and human biopsies. Bulk human RNA as well as spatial analysis of perifollicular T cells confirmed enrichment of *CD69* and *SELL/CD62L*. Different pathways predominated in other CD3+ regions of interest in CD4+ driven conditions including mucocutaneous lupus erythematosus and subacute cutaneous lupus erythematosus. Last, we identify novel drug-targetable pathways, namely *CFD* and *S100A8/9*, that could be further explored to disrupt processes in these conditions through veterinary and human trials.

## Introduction

Primary cicatricial alopecias (PCA) encompass several rare and common entities that are all characterized by hair loss, collagen deposition, and wound-healing type-responses in the absence of obvious physical injury^1^. These disorders can cause psychological stress and disfigurement^2^. The initiating factors for distinct clinical entities are likely different and include age/sex hormones^3^ in addition to genetics and environmental triggers^4^ that lead to collapse of immune privilege in the hair follicle^5^. Since most patients present to the clinic after disease onset, stopping inflammation and reversing the fibrotic process are of utmost importance for improving prognosis and chances of disease reversal. Furthermore, studying shared pathological processes in different disorders and models will yield better insights into fibrotic processes and may accelerate treatments for diseases with similar features.

The clinical subtypes of lymphocytic PCA include chronic cutaneous lupus erythematosus (CCLE, most common subtype of which is discoid lupus erythematosus, DLE)^6^, lichen planopilaris (LPP) and frontal fibrosing alopecia (FFA)^7^, among others^7^. The pathophysiology of these conditions shares similar features: inflammation, hair loss, and collagen deposition in the follicle. Different clinical subtypes of PCA have unique gene signatures: for example, CCLE has an interface dermatitis histological pattern in addition to hair follicle involvement, LPP appears to be more inflammatory (with higher numbers of CD8+ T cells^8^ and macrophages^9^), and FFA appears more fibrotic (with higher markers of epithelial to mesenchymal transition^9^).

A paucity of animal models has hindered preclinical studies of potential treatments of PCA. Here we present a novel mouse model that shares gene expression with several PCA entities that may prove useful for mechanistic studies of lymphocytic PCAs. Companion animals spontaneously develop many of the same diseases as their human companions and may also serve as large animal models of disease. For example, pet dogs can develop DLE and other forms of alopecia^10^. Dogs share the same environment and lifestyle as their human companions, and dogs owned by SLE patients have a higher risk of developing lupus themselves^11^. Previous work from us and others has characterized the transcriptome of DLE and demonstrated shared gene expression in bulk microarray samples ^12^. We sought to examine cellular neighborhoods and better define the microanatomical niche of T cells in and around hair follicles using spatial transcriptomics, and to catalog potential drug targets that could be tested in future veterinary and human clinical trials.

## Results

### Development of a mouse model of lymphocytic primary cicatricial alopecia that shares features and gene expression patterns with human lymphocytic PCA

Our lab has generated a mouse model that involves adoptive transfer of ovalbumin-specific CD8+ T cells (OT1) into recipient mice expressing ovalbumin in Keratin 5+ cells, which includes hair follicle keratinocytes (**Fig 1A**). This model is similar to the Katz lab model of GVHD^13^, but differs in that our recipients express the K5-ovalbumin under a tetracycline response element so that we can induce flares^14^. Our recipients are also TLR9-/-, which forces signaling through TLR7 as a pro-autoimmune signal ^15,16^. The resulting disease in our mice resembles PCA, including alopecia, fibrosis and mucin deposition (**Fig 1B-C**). Though this model is similar to GVHD models in skin disease score (**Fig 1D**), mice do not significantly lose weight compared to littermate controls (**Fig 1E**). Only rtTA+ K5TGO+ TLR9-/- recipients develop anti-nuclear antibodies (ANAs, **Fig 1F**) and splenomegaly (**Fig 1G**), not littermate controls.

**Figure 1.**
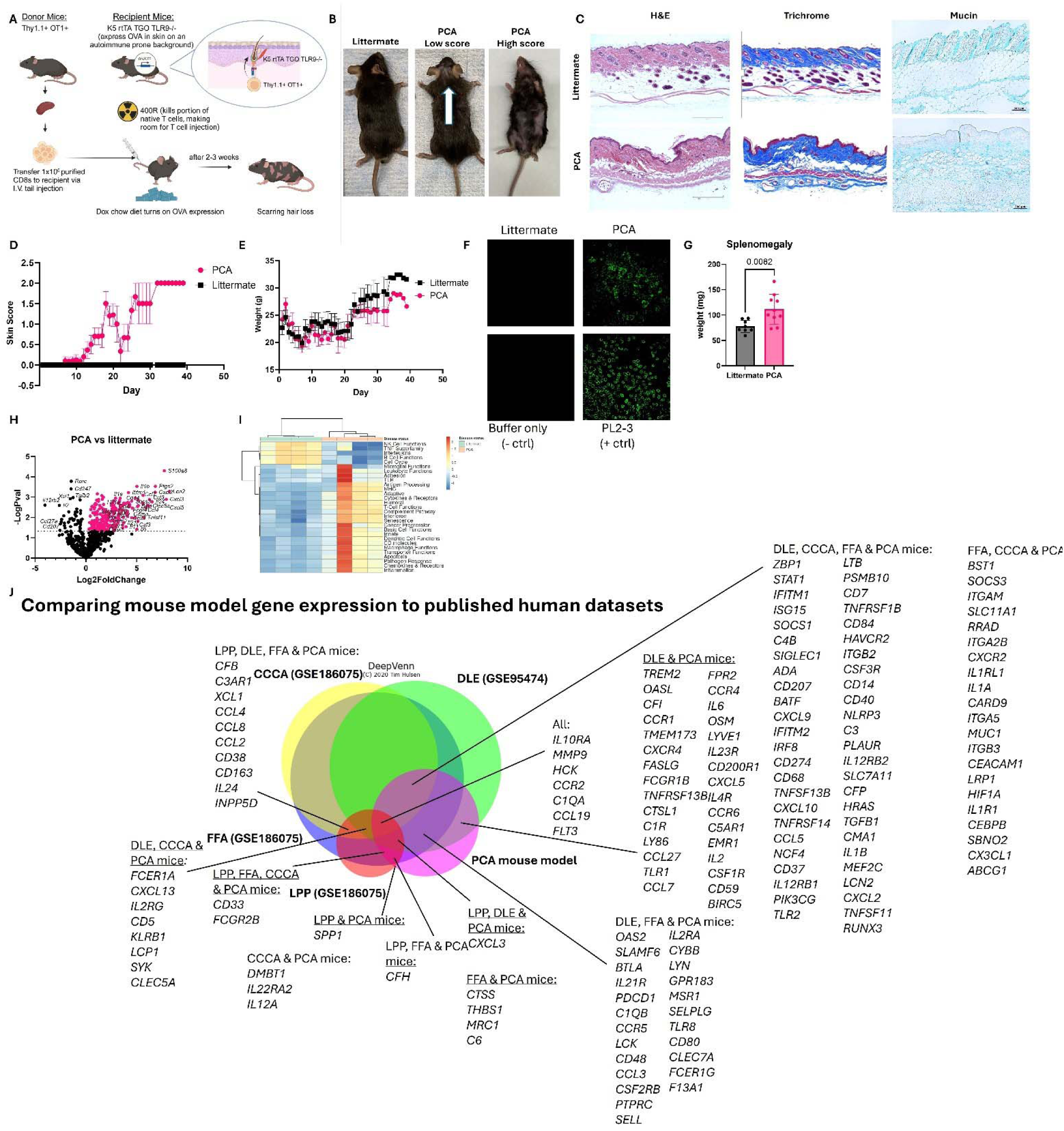
A mouse model of primary cicatricial alopecia (PCA) is dependent on antigen-specific CD8+ T cells. A. BioRender schematic of the PCA mouse model. B. Example photographs demonstrating hair loss in recipient mice. C. Example H&E, trichrome and mucin staining of skin sections from mice. D. Skin scores over time. E. Mouse weights over time. F. Example ANA staining. G. Spleen weights demonstrating splenomegaly in recipient mice (panels A-G n=8 littermate and 10 PCA mice compiled from 3-4 separate experiments). H. Microarray gene expression analysis in PCA mice versus littermate controls (n=4 per group). I. Gene set enrichment analysis of microarray data from mice. J. Comparison of DEGs from the PCA mouse model to publicly available human datasets.

We sought to characterize gene expression in lesional skin of PCA mice versus littermate controls using a microarray (**Fig 1H**). We noted significant increases in chemokines/cytokines, complement family members and CD markers (p<0.05, log2fold change +/-1.5). Gene set enrichment analysis revealed multiple inflammation-related pathways are upregulated in PCA mice versus littermate controls (**Fig 1I**). We compared our mouse model DEG list to published human datasets (GSE186075^17^ and GSE95474^6^). We find overlapping gene expression in our model as compared to human lymphocytic PCAs (**Fig 1J**), which makes sense given the model is dependent upon adoptive transfer of T cells. This is clinically relevant to human PCA, given TLR7 dysfunction in both alopecia^18,19^ and lupus erythematosus^20^, of which discoid lupus is one of the lymphocytic PCA clinical entities.

### Cytotoxic T cells interact with hair follicles and mediate destruction of CD34+ bulge cells in the PCA mouse model

Next, we asked what target cells the T cells were attacking in the PCA mouse model (**Fig 2A**). We performed flow cytometry on lesional skin from mice using a panel for flow cytometry of hair follicles^21^ (**Fig S1**). We noted significant decreases in CD34+ bulge cells from hair follicles and interfollicular epithelial cells in PCA mice compared to littermate controls (**Fig 2B-C**). Flow cytometry analysis of T cells from PCA mice indicated generation of both resident memory T cells (Trm) and recirculating memory T cells (Trcm) (**Fig 2D, Fig S2**). Specifically, we noted populations of T cells that express CD62L/L-selectin (**Fig 2E-F**), which is a binding partner of CD34^22^ and a marker of recirculating memory T cell populations^23^. We also noted significant increases in CD122 expression, which is part of the IL-15 receptor. Notably, IL-15 blockade prevents alopecia areata in C3H mice^24^. To observe T cell behavior in the follicles, we bred OT1 mice to Kikume photoconvertible mice and used cells from these donors to induce PCA^25^ (**Fig 2G**). We performed *in vivo* imaging of the ear skin^26^ in PCA mice and littermate controls 24h after photoconversion with a 405nm violet laser. We observed GFP+ cells moving quickly in littermate controls, likely in the vasculature (**Fig 2H, movie S1**). In contrast, we found RFP+ cells arrested near hair follicles, surrounded by clouds of GFP+ cells (**Fig 2I, movie S2**). Taken together, our data suggests that Trm and Trcm populations are recruited to and arrest at the hair follicle, are responsive to IL-15, and may target CD34+ cells through CD62L binding.

**Figure 2.**
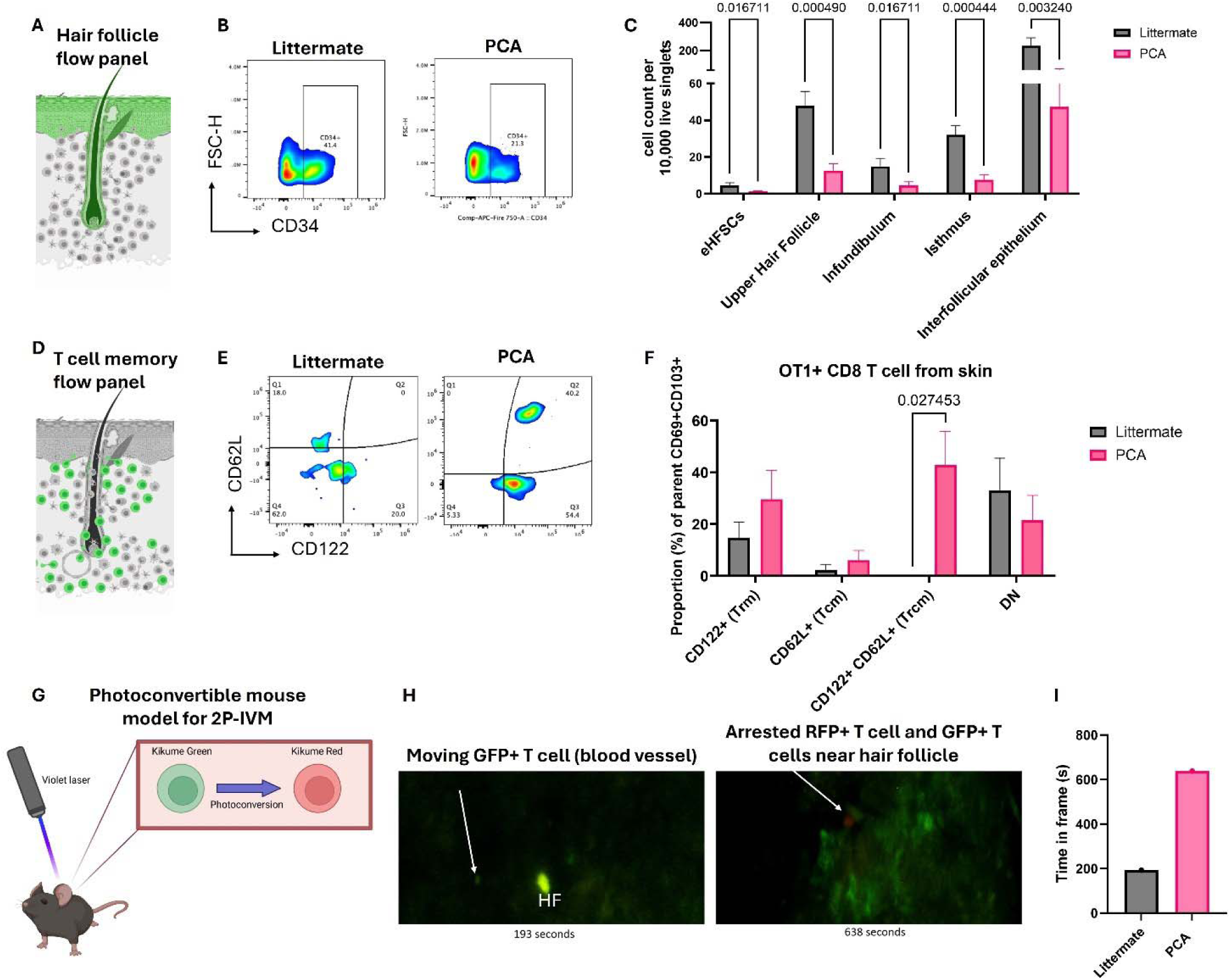
CD8+ T cells in the PCA mouse model express CD122 and CD62L, and arrest and attack CD34+ and other K5+ skin and hair cells. A. BioRender schematic of hair follicle flow panel. B. Example flow plots of CD34 expression in the skin of littermate versus PCA mice. C. Quantification of hair follicle cell types in littermates versus PCA mice (n=8 littermates and 12 PCA mice per group pooled from 4 experiments; multiple T tests with Bonferroni correction significant as indicated). D. Biorender schematic of T cell flow panel. E. Example flow plots of CD122 and CD62L expression. T cells were pregated on live, single, cell, CD45+CD3+Kikume green+/Thy1.1+CD69+CD103+ cells. F. Quantification of T cell phenotypes as a proportion of the parent gate. DN - double negative cells. G. BioRender schematic of photoconvertible mouse model for in vivo ear imaging. H. Example flow plots of Kikume green expressing T cells in the skin of recipient mice. I. Stills from 2 photon in vivo ear imaging videos. J. Quantification of time spent in frame by the T cells.

### Spatial transcriptomics of PCA mouse skin compared to CLE mouse skin reveals different pathways allowing for interactions between CD8 versus CD4 T cells and stromal cells

We wanted to further characterize potential binding interactions between T cells and hair follicle cells in our mouse model. We hypothesized that autoreactive CD8+ T cells use discreet recognition pathways for interacting with hair follicle stromal cells as compared to CD4+ T cells. Therefore, we performed transcriptomics on skin from CD8+ recipients (PCA mice), CD4+recipients (CLE mice) and littermate controls (**Fig 3A**). First, we analyzed total gene expression in skin tissue using a bulk microarray (**Fig 3B**). We noted unique and overlapping genes depending upon whether the rtTA+ K5TGO+ TLR9-/- recipients received OT2 (CD4+) or OT1 (CD8+) T cells (**Fig 3C**). We used Digital Spatial Profiling with CD3, CD8, and CD45 morphology markers to identify T cells and hair follicles (**Fig 3D**). We used cell segmentation approaches to pull out CD3+CD8+, CD3+CD8- and other CD45+ cells from regions of interest (ROIs), and used geometric ROIs to pull out hair follicles and keratinocytes for comparators (**Fig 3E, Fig S3**). Comparing gene expression in the hair follicle from PCA vs healthy mice (**Fig 3F**) and CLE vs healthy mice (**Fig 3G**) revealed many differentially expressed genes. Examples of genes that were upregulated in the hair follicle from both mouse models include *ATF4*, a transcription factor that orchestrates the integrated stress response in hair follicles^27^ (**Fig 3H**), *HMGB1*, a danger signal that induces hair regrowth following trauma^28^ but serves as a key danger-associated molecular pattern in lupus^29^ (**Fig 3I**), and *CXCL16*, a chemokine that is important for generation of Trm in the context of melanoma^30,31^ (**Fig 3J**). *CD200R1*, a classical immune privilege signal in the hair follicle^32^, was reduced in both models (**Fig 3K**).

**Figure 3.**
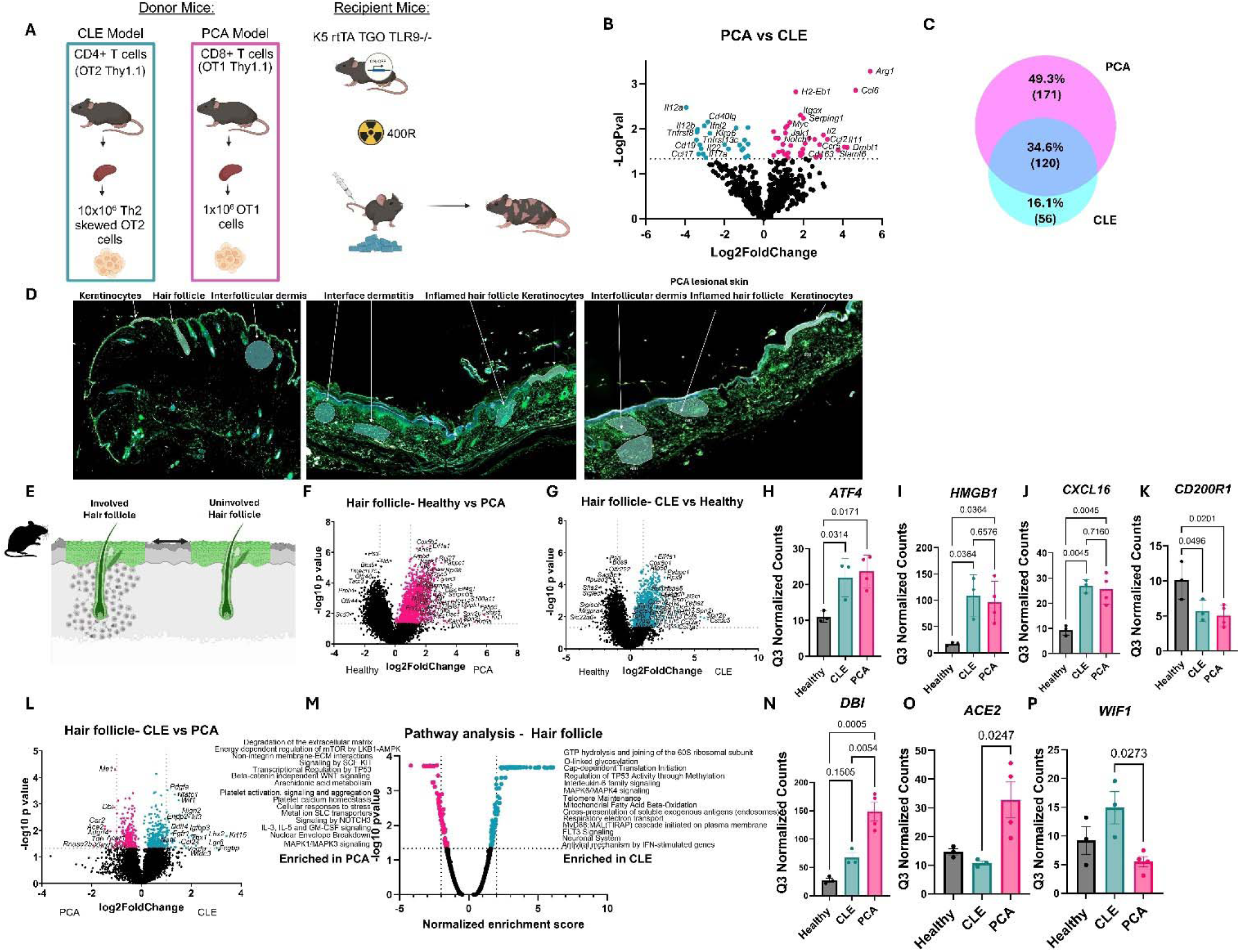
Spatial transcriptomics of PCA mouse model CD8 recipient mice compared to CD4 recipient mice reveals conserved and differential gene expression. A. BioRender schematic demonstrating CD4 (CLE) versus CD8 (PCA) recipient mouse models. B. Bulk microarray data from CLE versus PCA mouse skin. C. Biovenn analysis of unique and shared DEGs from the skin of recipient mice. D. Example images for ROI selection. E. BioRender schematic of involved versus uninvolved/healthy hair follicle comparison. F. PCA versus healthy and G. CLE versus healthy hair follicle gene expression. H. *ATF4*, I. *HMGB1*, and J. *CXCL16* were upregulated in both CLE and PCA hair follicles compared to healthy. K. *CD200R1* expression was lost in hair follicles in both models. L. Comparing PCA versus CLE hair follicle gene expression. M. Pathway analysis of PCA versus CLE gene expression. N. *DBI* and O. *ACE2* were upregulated in PCA hair follicles compared to CLE. P. *WIF1* expression was lower in PCA hair follicles compared to CLE. (n=4 healthy, 4 CLE and 4 PCA tissues sectioned; ROIs analyzed that passed QC include 3 healthy, 3 CLE and 4 PCA hair follicles).

We compared hair follicles from CLE vs PCA mice, and found many DEGs (**Fig 3L**). Pathway analysis of these genes demonstrated terms including “degradation of the extracellular matrix (ECM)”, “non-integrin membrane-ECM interactions”, and “platelet activation, signaling and aggregation” specifically in PCA mouse hair follicles (**Fig 3M**). Notably, *DBI* (**Fig 3N**), which is involved in lipid metabolism, and *ACE2* (**Fig 3O**), which is the major receptor associated with Sars-CoV-2 cell entry and has been implicated in androgenic alopecia^33^ and post-COVID-19 disease ^34^ or vaccination-associated alopecia^35^ were significantly upregulated in PCA hair follicles. In contrast, *WIF1* (**Fig 3P**), the loss of which is associated with baldness^36^, was downregulated compared to CLE mice. Taken together, these data demonstrate unique and overlapping hair follicle danger signals in the B6 mouse models, and show that CD4 versus CD8 autoreactivity may result in slightly different responses in hair follicles, with CD8 specifically contributing to ECM reorganization and potentially hair follicle fibrosis.

### Examination of spatial gene expression in spontaneous discoid lupus erythematosus and mucocutaneous lupus in client-owned dogs reveals pathways conserved with mouse and human PCA

Dogs also spontaneously develop DLE^10^. Our lab previously characterized bulk gene expression in lesional skin from companion dogs that spontaneously developed DLE using a microarray^12^. Here, we compared healthy leg margin skin excisions from amputations to DLE and mucocutaneous lupus erythematosus (MLE) biopsies^37^ (**Fig 4A**). First, we performed bulk RNA sequencing on curls from FFPE blocks from DLE cases versus healthy margins (**Fig 4B**). We noted a loss of *CD34* in DLE compared to healthy margins, and upregulation of chemokines (*CCL5*), keratins (*KRT3, 4, 13, 76, 222*), interferon response genes (*MX1, PSMB8*), and S100 family genes (*S100A8*).

**Figure 4.**
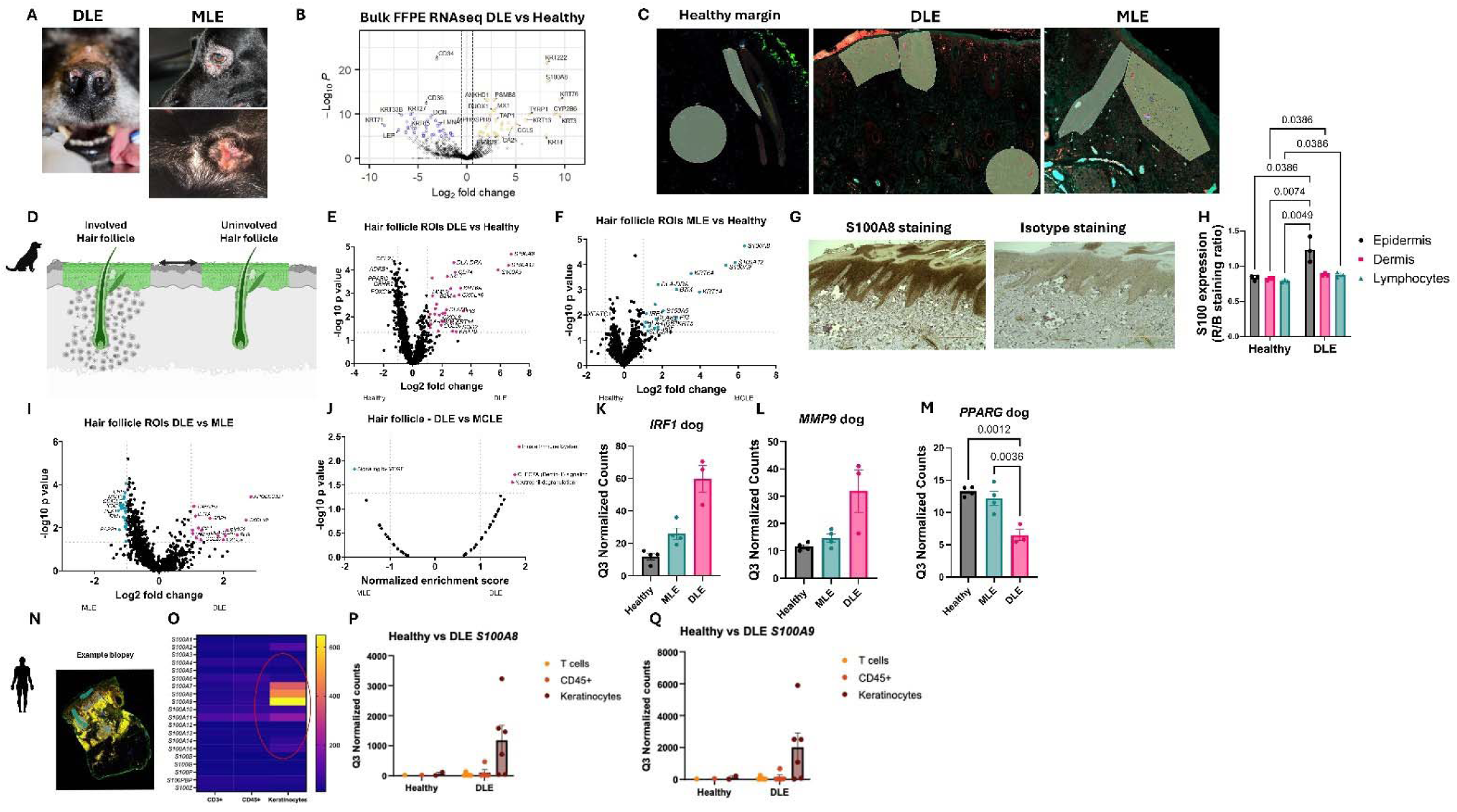
Spatial transcriptomics of spontaneous discoid (DLE) and mucocutaneous lupus (MLE) in client owned dogs demonstrates increased S100 expression and loss of PPARG as pertinent to human discoid lupus. A. Left: example clinical photos of canine facial DLE featuring scarring, dyspigmentation, erosions/ulcerations and loss of nasal planum architecture. Right: example clinical photos of periorbital (top) and genital (bottom) MCLE in dogs. B. Bulk RNAseq from DLE and healthy margin FFPE curls demonstrates upregulation of chemokines, keratins and S100 genes. C. Example ROI selection from canine sections. D. BioRender schematic for involved versus uninvolved hair follicle comparisons. E. DLE versus healthy and F. MLE versus healthy hair follicle comparisons. G. Example S100A8 staining compared to isotype staining. H. Quantification of S100A8 staining in different microanatomical compartments determined by ImageJ. I. Comparison of DLE versus MLE hair follicle gene expression. J. Pathway analysis revealed innate immune system, CLEC7A signaling and neutrophil degranulation were enriched in canine DLE hair follicles. K. *IRF1* and L. *MMP9* were upregulated specifically in DLE hair follicles. M. DLE hair follicles revealed a significant loss of *PPARG* compared to MLE and healthy. N. Example human DLE biopsy (GSE182825). O. Heatmap of S100 family gene expression divided by ROI. *S100A8/9* were among the top upregulated genes in keratinocytes. P. *S100A8* and Q. *S100A9* quantification in individual ROIs reveals significant increases in DLE keratinocytes.

We performed Digital Spatial Profiling with the Canine Cancer Atlas (CCA) probe set using CD3, CD8 and CD45 morphology markers. We employed a similar ROI selection as in the mouse WTA, selecting hair follicles as well as T cell and immune cell ROIs (**Fig 4C**), and examined involved versus healthy hair follicles from our spatial data (**Fig 4D**). Comparing gene expression in the hair follicle from DLE versus healthy skin (**Fig 4E**) and MLE vs healthy skin (**Fig 4F**) revealed many differentially expressed genes. Some of the highest upregulated genes were S100 family genes. We confirmed expression of S100A8 at the protein level in keratinocytes using immunohistochemistry (IHC) (**Fig 4G**), and found that staining was enriched in the epidermis and hair follicles of DLE compared to the dermis and lymphocyte-rich regions and to healthy skin (**Fig 4H**).

We compared hair follicles from DLE vs MLE dogs and found many DEGs (**Fig 4I**). “Signaling by VEGF” pathway was enriched in MLE hair follicles, whereas “Innate Immune System”, “CLEC7A (Dectin-1) signaling” and “Neutrophil degranulation” were enriched in DLE hair follicles (**Fig 4J**). *IRF1*, which is an interferon response gene previously reported to be upregulated in plucked human hair follicles from chronic DLE patients^38^, and *MMP9*, a matrix metalloproteinase previously reported to be elevated in human CCCA^39^, were significantly upregulated in DLE dog hair follicles (**Fig 4K-L**). There was also a significant loss of *PPARG*, which when deleted from hair follicle stem cells causes scarring alopecia in mice^40^ (**Fig 4M**).

We queried keratinocyte ROIs from our previously deposited human healthy, DLE and SCLE WTA dataset GSE182825^41^ to determine if S100 family genes are conserved across canine and human (**Fig 4N**). Comparing CD3+ CD45+ and CD45-keratinocyte geometric ROIs demonstrated highest counts of *S100A7, A8* and *A9* in keratinocytes (**Fig 4O**). We quantified *S100A8* and *A9* (**Fig 4P & Q**) given that these were also elevated in the canine dataset and found that indeed they are upregulated in keratinocyte ROIs compared to CD3+ and CD45+ ROIs. Taken together, our comparative spatial transcriptomics of canine DLE demonstrates conservation of S100 family proteins in keratinocytes, as well as IFN responses and loss of protective factors in hair follicles including *PPARG*, which is also downregulated in human LPP^42^.

We compared perifollicular versus interfollicular CD8+ T cells in our mouse model and spontaneous DLE in pet dogs. Several DEGs overlapped, including *CXCL12* (**Fig S4**). Pathway analysis using the GeoMX pathways tool identified “hemostasis” and “platelet activation” terms as conserved. This is interesting, given that platelet rich plasma (PRP) is used as a treatment for PCA in people^43^. Taken together, these data suggest that CD8+ T cells in hair follicles are capable of producing CXCL12, and express genes that may relate to the mechanism of action of PRP therapy.

### Human discoid lupus erythematosus biopsies exhibit markers of T cell memory, chemokine ligands and fibrotic responses in and around hair follicles

DLE disproportionately affects skin of color patients^44^. Therefore, we sought to perform spatial transcriptomics on DLE biopsies from archival blocks from skin of color patients to better understand the microanatomical niche in this patient population (**Fig 5A**). First, we performed bulk RNA analysis using a microarray (**Fig 5B**). We noted significant upregulation of markers identified in our mouse model (*CD69, SELL, IL2RB/CD122*), chemokines (*CCL3, CXCL9, CXCL11*), cytokines (*TNF*), IFN response genes (*IFIT1, MX1, MX2, IRF7, ISG15, STAT1*), as well as markers previously associated with lupus (*MYD88, FASLG*). Cell type enrichment analysis revealed significant increases in cytotoxic cells, B cells, neutrophils and T cells compared to healthy margin skin (**Fig 5C**).

**Figure 5.**
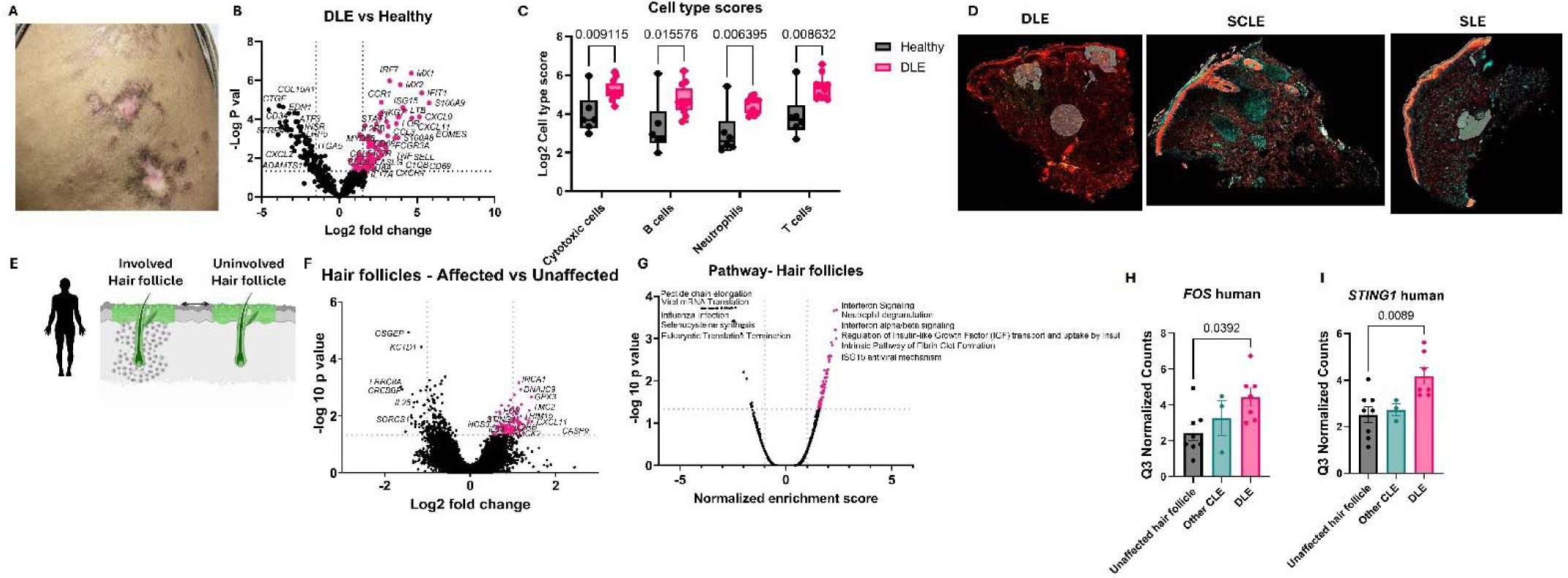
Spatial transcriptomics of DLE biopsies from skin of color patients reveals *FOS* and *STING1* expression in affected hair follicles. A. Example clinical photo of DLE. B. Bulk microarray data from matched biopsies reveals upregulation of *CD69, SELL*, chemokines and IFN response genes. C. Cell type score quantification reveals increases in cytotoxic cells, B cells, neutrophils and T cells compared to healthy margin skin (n= 6 healthy and 12 biopsies from 10 DLE patients). D. Example ROI selections from biopsies. E. BioRender schematic of affected vs unaffected hair follicle comparison. F. Gene expression in DLE affected versus unaffected hair follicles. G. Pathway analysis reveals interferon signaling, neutrophil degranulation, IFN alpha/beta signaling, regulation of IGF, intrinsic pathway of fibrin clot formation and ISG15 antiviral mechanism as top pathways enriched in DLE hair follicles. H. *FOS* and I. *STING1* were significantly upregulated in affected hair follicles. (ROIs from n=9 acute DLE, 10 DLE, 13 SLE, 3 SCLE and 12 tumid LE).

We used the Whole Transcriptome Atlas (WTA) to compare DLE blocks to other types of non-scarring CLE, including SCLE and SLE (**Fig 5D, Table S3**). We examined affected hair follicles from DLE and acute DLE patients compared to unaffected hair follicles from all CLE biopsies (**Fig 5E**). Affected follicles specifically upregulated *CXCL11, ROCK2* and *CASP9* among other genes (**Fig 5F**). Pathway analysis revealed “interferon signaling”, “neutrophil degranulation”, “interferon alpha/beta signaling”, “regulation of IGF transport and uptake”, “intrinsic pathway of fibrin clot formation” and “ISG15 antiviral mechanism” terms were enriched in DLE hair follicles (**Fig 5G**). Interestingly, *FOS*, which is involved in wound healing^45^, and *STING1*, which is involved in hair follicle stem cell replicative stress^46^, were significantly upregulated in affected DLE hair follicles (**Fig 5H-I**). Taken together, our data confirm previously reported lupus-associated gene expression and identify novel pathways that have not been studied in human DLE.

### T cell-hair follicle gene ontology terms “metabolic process”, “localization” and “response to stimulus” are conserved across mammalian species

We compared T cells in hair follicles to interfollicular T cells to better understand how they interact with T cells during primary cicatricial alopecia. We pooled all T cell ROIs and analyzed perifollicular vs interfollicular ROIs (**Fig 6A**). In PCA mice, *CXCL12, CLIP, ZFP146, MET, OPN5, SCL35D2* were enriched in perifollicular T cells, whereas *SOCS3, SERPINB13*, and *IRX3* were enriched in interfollicular T cells (**Fig 6B**). We exported the DEG lists and, using p<0.05 and log2FC cutoff of 1.0, we used PantherDB to analyze Gene Ontology (GO) pathways as Biological processes. The top signatures excluding non-annotated genes were “cellular process”, “biological regulation”, “metabolic process”, “localization” and “response to stimulus” (**Fig 6C**). In DLE dogs, *CXCL12* was conserved in perifollicular T cells (**Fig 6D**). We also noted *IFNG*, which has previously been described as a “node” in DLE^47^, was upregulated in perifollicular T cells whereas *FOS* was upregulated in interfollicular T cells. GO pathways were similar, with “cellular process”, “biological regulation”, “metabolic process”, and “response to stimulus” being enriched in perifollicular T cells (**Fig 6E**). In human DLE samples, we noted *CD69* and *SELL* were enriched in perifollicular T cells, along with *IFNL3, CCL3, CCL8* and *BACH1* genes (**Fig 6F**). Interfollicular T cells were enriched for expression of *MAP2K5, SFRP2, CCL17* and *GATA5* (**Fig 6F**). GO pathways were again similar, with “cellular process”, “biological regulation”, “metabolic process”, “localization” and “response to stimulus” as being the most enriched upregulated terms (**Fig 6G**). Taken together, these data suggest that T cell metabolism and migration are key biological processes conserved across species in PCA.

**Figure 6.**
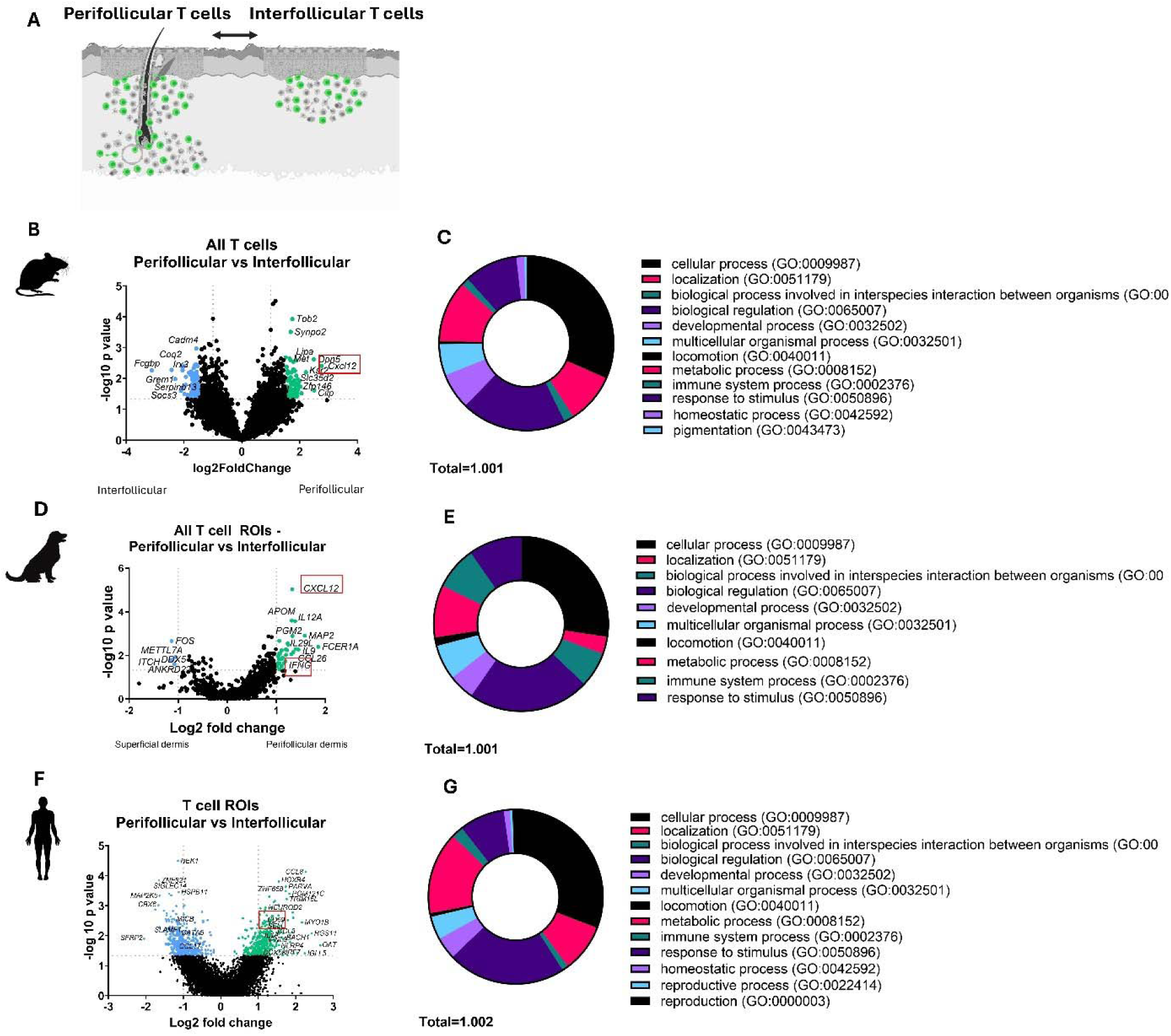
Comparative spatial transcriptomics of mouse, dog and human perifollicular versus interfollicular T cells reveals key pathways including localization and metabolism. A. BioRender schematic of perifollicular vs interfollicular T cell gene expression comparison. B. Mouse peri versus interfollicular T cell gene expression. C. Gene ontology analysis of DEGs with P<0.05 and log2 fold change 1+ using PantherDB reveals specific biological processes including metabolism and localization processes. D. Dog peri versus interfollicular T cell gene expression. E. Gene ontology analysis of DEGs with P<0.05 and log2 fold change 1+ using PantherDB reveals specific biological processes including metabolism and localization processes. F. Human peri versus interfollicular T cell gene expression comparison. G. Gene ontology analysis of DEGs with P<0.05 and log2 fold change 1+ using PantherDB reveals specific biological processes including metabolism and localization processes.

We then wanted to compare hair follicles themselves across species (**Fig 7A**). We again exported the DEG lists and, using p<0.05 and log2FC cutoff of 1.0, we used PantherDB to analyze GO pathways as Biological processes. The top signatures in mouse PCA excluding non-annotated genes were “cellular process”, “biological regulation”, “metabolic process”, “localization” and “response to stimulus” (**Fig 7B**). Based on our previous individual species analyses, we noted CXCR3 ligands were upregulated across species. *CXCL9* was significantly upregulated in PCA hair follicles (**Fig 7C**). We also examined IFN response genes. *IRF1* was upregulated in both CLE and PCA mouse models (**Fig 7D**). Last, we examined spatial *CFD* expression, which was a top DEG in bulk microarray data. *CFD* was highest in deep dermal ROIs (**Fig 7E**). In dog affected hair follicles, the top GO terms were “cellular process”, “biological regulation”, “response to stimulus”, “metabolic process” as well as “developmental process” and “multicellular organismal process” (**Fig 7F**). *CXCL10* was the key CXCR3 ligand upregulated in affected hair follicles (**Fig 7G**), and *IRF1* was conserved as a key IFN response gene (**Fig 7H**). *CFD*, however, was not enriched in affected hair follicles, but was significantly higher in perifollicular T cells (**Fig 7I**). In human DLE affected hair follicles, “cellular process” “biological regulation”, “response to stimulus” “metabolic process” as well as “developmental process” and “multicellular organismal process” were the key GO terms (**Fig 7J**). *CXCL11* was the key CXCR3 ligand upregulated in affected hair follicles (**Fig 7K**). *IRF1* was not significantly higher in DLE affected hair follicles; rather *IFIT1* was significantly upregulated in hair follicles (**Fig 7L**). *CFD* was enriched in CD45+ ROIs (**Fig 7M**). Taken together, these data suggest that for proper understanding of biological processes across species, it is necessary to also have knowledge of the functional gene systems to know which genes can compensate for one another in a given biological system, such as the CXCR3 ligands. Further, similar gene expression networks might be expressed by different cell types, as exemplified by *CFD*. Our data also recapture the CXCR3 and IFN pathways as key biological processes that are conserved across mammalian species during PCA.

**Figure 7.**
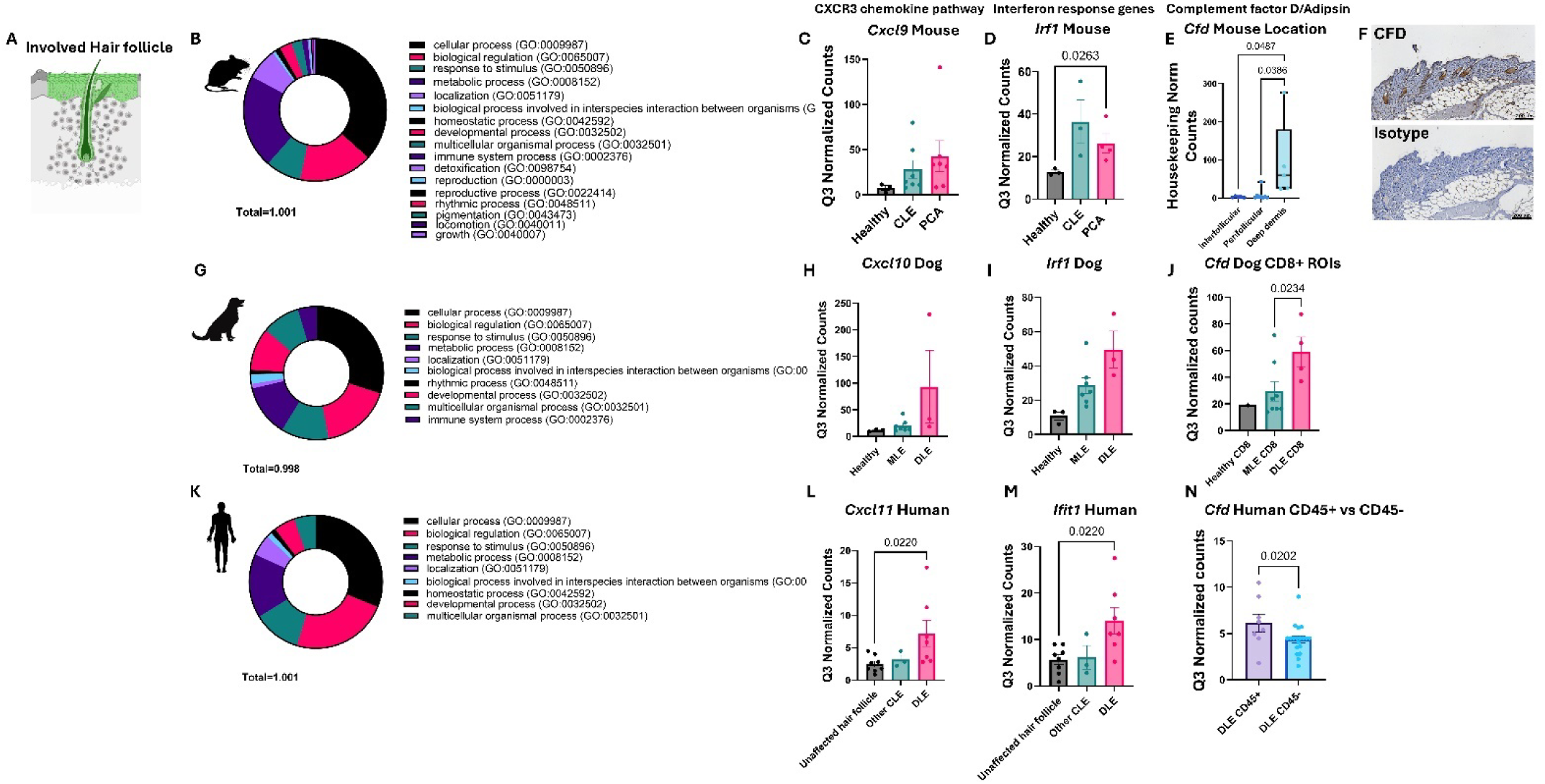
Comparative spatial transcriptomics of mouse, dog and human hair follicles reveals conserved pathways and different cellular contexts of key DEGs. A. BioRender schematic of involved hair follicle cross species comparison. B. Mouse gene ontology processes in hair follicles as assessed by positive DEGs. C. *CXCL9* and D. *IRF1* expression are increased in PCA hair follicles. E. *CFD* is increased specifically in the deep dermis of mouse skin. IHC inset confirms protein level expression of CFD. F. Dog gene ontology processes in hair follicles as assessed by positive DEGs. G. *CXCL10* and H. *IRF1* expression are increased in DLE hair follicles. I. *CFD* is increased specifically in CD8+ T cell ROIs. J. Human gene ontology processes in hair follicles as assessed by positive DEGs. K. *CXCL11* is increased in hair follicles. L. *IFIT1* but not *IRF1* is specifically upregulated in hair follicles. M. *CFD* is enriched in human CD45+ ROIs compared to CD45-ROIs.

## Discussion

Here, we present a comparative spatial transcriptomics study of hair follicles and T cells in the context of PCA. Our data recapitulate previous studies identifying key pathways and processes in PCA, such as IFN^47^, sterol biosynthesis^48^ and fibrosis^17^. We also identify potential novel targets. *CFD* is a highly upregulated DEG in our novel PCA mouse model, which is expressed in mouse hair follicles, dog T cells and human immune cells. While the cellular context of this signal was slightly different, the complement pathway terms were identified in several datasets presented here, and C3 immunofluorescence staining occurs in hair follicles in CCCA biopsies^49^. Recent studies have demonstrated a link between CD8+ T cells and complement system activation, showing that GZMK-expression CD8+ T cells activate complements and promote inflammation by cleaving complements such as C2, C3, C4 and C5^50^. This evidence supports our findings that complement pathways are relevant in PCA, further emphasizing the potential target of complement pathways. The CFD inhibitor Danicopan is currently FDA-approved for extravascular haemolysis (EVH) in adults with paroxysmal nocturnal haemoglobinuria (PNH)^51^. Danicopan is currently being tested in clinical trials for Geographic atrophy (GA), a severe form of age-related macular degeneration (AMD)^52^. It would be interesting to test drugs in the complement inhibitor class such as C1, C3, C5 and Factor B or D inhibitors for topical, intralesional and, if necessary, oral administration preclinically as well as in veterinary and human trials for PCA, especially given that they eye and hair follicle are both immune privileged organs.

Similar to human DLE, canine DLE is responsive to hydroxychloroquine^53^, tacrolimus^54^ and was recently shown to be responsive to the JAK inhibitor oclacitinib^55^. Here we show that S100A8/9 were highly conserved between canine and human in DLE keratinocytes. Small molecule inhibitors of S100A8/9 are in preclinical development. Most studies have used these inhibitors to treat myocardial dysfunction^56^, with some recent work demonstrating S100A8/9 inhibition prevents COVID-19 induced lung damage^57^. Additionally, these proteins have been associated with liver fibrosis associated with cirrhosis^58^. It would be interesting to test drugs in this class for topical, intralesional and, if necessary, oral administration in veterinary and human trials.

Adachi et al demonstrated that hair follicle-derived IL-7 and IL-15 mediate skin-resident memory T cells at homeostasis and during cutaneous T cell lymphoma (CTCL), specifically demonstrating dependency of CD8+ Trm on IL-15 and IL-7, and CD4+ Trm on IL-7^59^. Christiano lab demonstrated that either IL7 or IL15 blockade prevents alopecia areata in C3H mice^24,60^. Here we show increased CD122 expression and IL7 signaling pathways are recapitulated in our B6 PCA mouse model. However, IL15 can protect hair follicle immune privilege in *ex vivo* hair follicle cultures from Alopecia Areata patients^61^, promoting proliferation of stem cells. Further investigation is required to understand the context of IL15 signaling in scarring alopecia. We also noted CD69+ CD62L+ (*SELL*) T cells near hair follicles. These cells have previously been called recirculating memory T cells (Trcm)^23,62^. Given that the RFP+ T cells also recruited GFP+ T cells as determined by 2 photon microscopy, it is likely that the GFP+ T cells are responsible for attacking CD34+ bulge stem cells through CD62L tethering. Notably, CD62L is being developed as a target for the treatment of chronic lymphocytic leukemia (CLL)^63^. Further investigation is warranted to determine how to disrupt this process to maintain hair follicle stem cells in PCA.

Limitations of our study include small sample size, targeted panels and limited morphology markers. We also recognize one epidemiological study that did not find a link between TLR7 SNPs and DLE in a Polish population^64^. Despite this, higher TLR9 expression in cutaneous lupus is associated with positive response to hydroxychloroquine^65^, and given that our mouse model still recapitulates gene expression in human DLE, we may be able to use this for exploration of novel CLE treatments for recalcitrant patients. DBI total body knockout mice exhibit alopecia^66^. However, in our mouse model, we noted an increase in DBI expression in PCA hair follicles. Further studies would need to be conducted to better understand the context of DBI activity in the TLR9KO lupus-prone background. Last, our human biopsies were aged, resulting in lower counts, though samples did pass quality control. In the future, we plan to prospectively collect samples for flash frozen RNA spatial analyses, and to employ spatial sequencing tools in development such as the 10X Genomics canine panel.

## Methods

### Mouse model

Mouse studies were conducted at UMass Chan on an IACUC-approved protocol (#202100229) at an AAALAC approved facility. We generated a B6 version of the previously published Balb/c cutaneous lupus erythematosus (CLE) model^67^ by breeding K5-TGO mice^14^ to TLR9-/- mice^68^ to generate lupus-prone recipients that express the OVA model autoantigen in keratin 5 expressing cells. To model CLE, we used OT2 T cells (JAX # 004194) instead of DO.11 T cells to induce disease in these mice by transferring 10 million Th2 skewed T cells as previously described^69^. To model PCA, we transferred 1 million freshly isolated OT1 T cells (JAX # 003831) to induce disease. This model is similar to the Okiyama and Katz OT1 transfer model of GVHD in WT recipients that includes inflammation and fibrosis of the skin and hair^13^; however our mice are TLR9KO which makes them lupus-prone. All T cells were isolated from donor spleens using MojoSort kits (Biolegend) per the manufacturer’s instructions.

### Canine biopsies

Biopsies from client-owned companion dogs were selected from the Tufts Cummings School of Veterinary Medicine biorepository. Samples were deposited with written owner consent on an IACUC-approved protocol. Cases were re-reviewed by a board-certified veterinary dermatologist (RMA) and were sectioned onto Leica bond plus slides after floating in RNAse free water for use in the Canine Cancer Atlas (CCA) Digital Spatial Profiling assay (NanoString).

### Canine FFPE Bulk RNAseq

RNA was isolated from 30uM curls using the Qiagen RNEasy FFPE kit per the manufacturer’s instructions. Samples were shipped to GeneWiz for processing and sequencing, using the NEBNext Ultra II RNA library prep kit with sequencing on Illumina HiSeq 4000 with paired-end 150 bp read configuration. FASTQ files were aligned to canfam 1 for further processing.

### Human biopsies

Human DLE scalp biopsies were selected from the Howard University biorepository on an archival tissue IRB protocols that were approved by the UMass Chan IRB and the Howard University IRB, with a Memorandum of Understanding (MOU) for cross-institutional studies. Cases were re-reviewed by a board-certified pathologist (BS) and dermatologist (CF). Blocks were sectioned onto Leica bond plus slides in the UMass Chan morphology core and were prepped for spatial analyses in the UMass Chan SCOPE core.

### Spatial transcriptomics

Biopsies from affected hair-bearing skin from mice, dogs or humans were sectioned onto Leica Bond Plus slides and were stored at 4C in slide boxes with desiccant until use (3 weeks maximum). NanoString GeoMX Digital Spatial Profiling platform was used for cross-species comparisons. Mouse and Human Whole Transcriptome Atlas (WTA) assays and Canine Cancer Transcriptome Atlas (CCA) were used to analyze samples. CD3, CD8 and CD45 were used as morphology markers with SYTO13 nuclear stain (antibodies in **Table S1**). GeoMX sample preparation was performed in the UMass Chan SCOPE core (RRID:SCR_022721).

Region of interest (ROI) selection was based on hair follicle morphology and cell segmentation using the same ROI selection methods for mice, dogs and humans for consistency. Collection plates were sequenced on an Illumina HiSeq in the UMass Chan High Throughput Sequencing Core. Data were analyzed in the GeoMX data analysis suite including biological probe QC, and Q3 normalization. For T cell ROIs, housekeeping gene normalization was also performed using the NanoString housekeeping gene list from the human TCR microarray panel.

### Comparison to publicly available datasets

We performed gene expression analysis comparison of skin from our mice to GEO datasets GSE186075^17^ and GSE95474^6^ from human PCA. Top tables were generated using Geo2R software and were exported into Excel. The human gene lists were truncated to match the common denominator genes in the mouse microarray panel as previously described using the VLOOKUP IS ERROR function^12^. Tables were sorted based on p value and normalized expression, and differentially expressed genes (DEGs) were compared using BioVenn software^70^.

### Histology

H&E staining, trichrome staining and IHC staining were performed in the UMass Chan Morphology Core. Complement Factor D polyclonal antibody (Thermo cat # PA579034) or isotype control were used at 1:100 dilution using Leica Bond Plus autostainer.

### Flow cytometry

Healthy mice skin and PCA mice skin were harvested, minced into small pieces, and digested in the skin digesting buffer with 2.0 mg/ml collagenase XI from Clostridium histolyticum (Sigma-Aldrich), 0.5 mg/ml hyaluronidase from bovine testes (Sigma-Aldrich), and 0.1 mg/ml DNAse (Sigma-Aldrich) for 40 minutes at 37°C. The single cell suspensions were washed with RPMI 1640 (Corning, #20623011), filtered through a 40 μm cell strainer (Fisher Scientific, #22363547), and stained with fluorescent antibodies (**Table S2**) for flow cytometry. The stained samples were analyzed using a Cytek Aurora cytometer (Cytek) and data analysis was performed with FlowJo 10.10.0 software.

### 2 photon in vivo microscopy

We performed in vivo imaging of ear skin in mice that received Kikume OT1 T cells as previously described^26^. Briefly, mice were anesthetized with ketamine/xylazine solution and were placed in a 3D printed stage. Images were captured with a Bliq photonics 2 photon microscope with 100nm averaging at 920nm. Images were compiled in Fiji and total duration of interactions of T cells with skin is reported.

### Data analysis & statistics

Two-way comparisons for differentially expressed genes (DEGs) and pathway analyses were performed in GeoMX software to calculate volcano plots, and individual hypotheses of gene families were tested by querying specific gene sets within the Q3 normalized count file. Data were graphed with GraphPad Prism software, and ANOVAs with post-tests were performed using p<0.05 as a cutoff.

## Supplemental Figures & Legends

**Figure S1.**
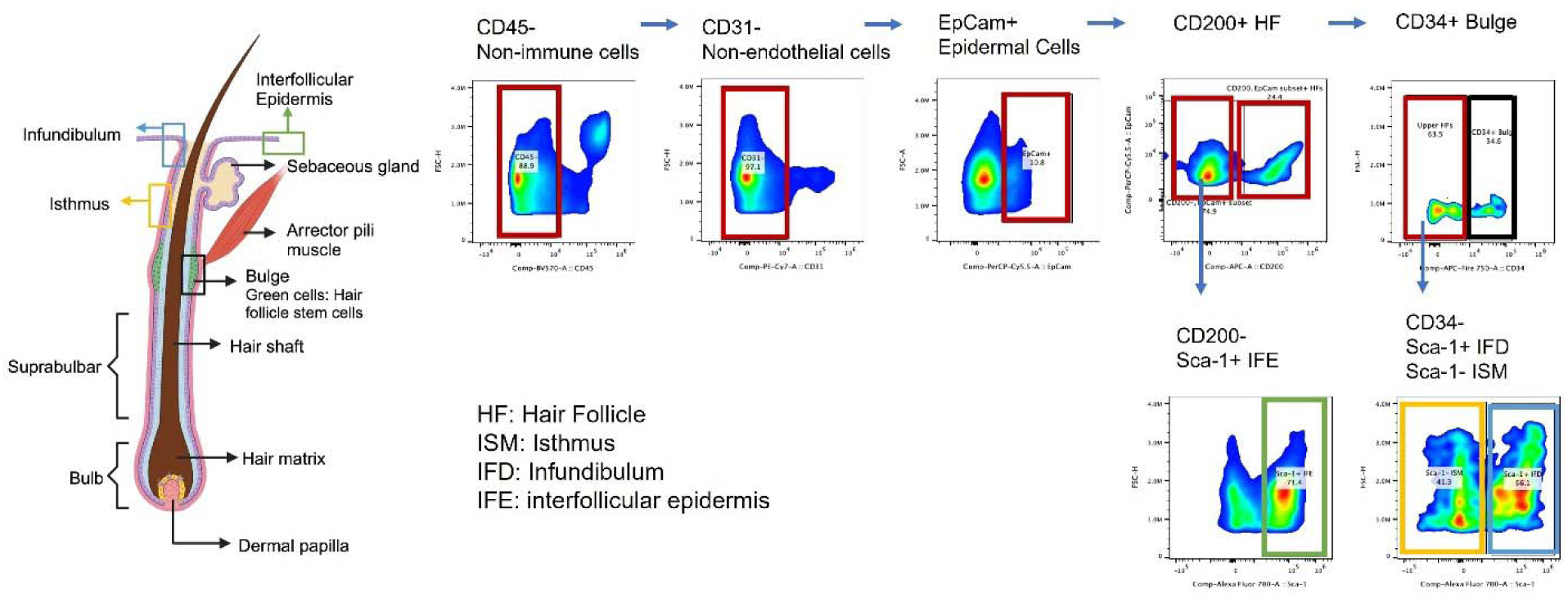
Example hair follicle flow cytometry gating. Colored boxes correspond to the cells and structures in the diagram (created with Biorender.com)

**Figure S2.**
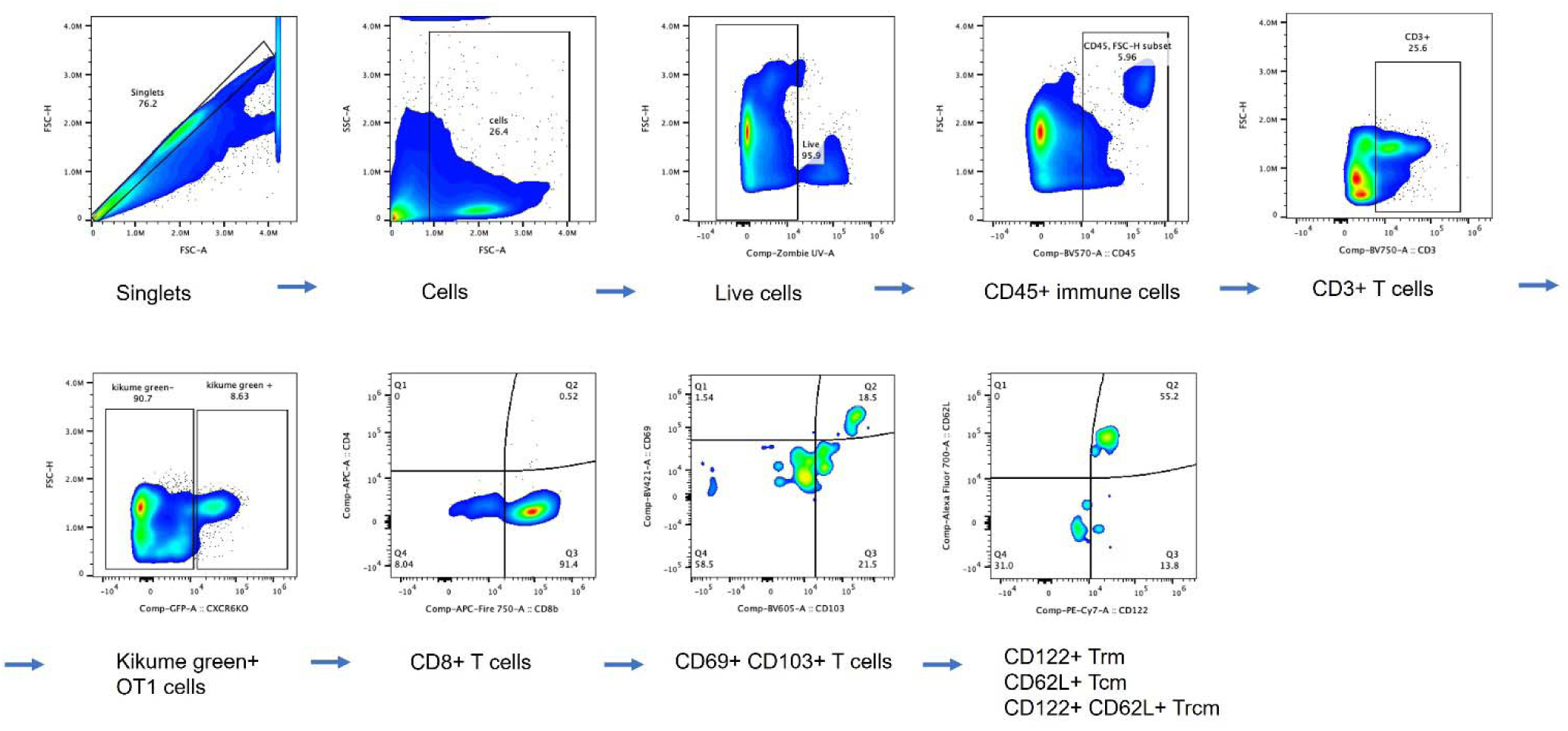
Example memory T cell flow cytometry gating.

**Figure S3.**
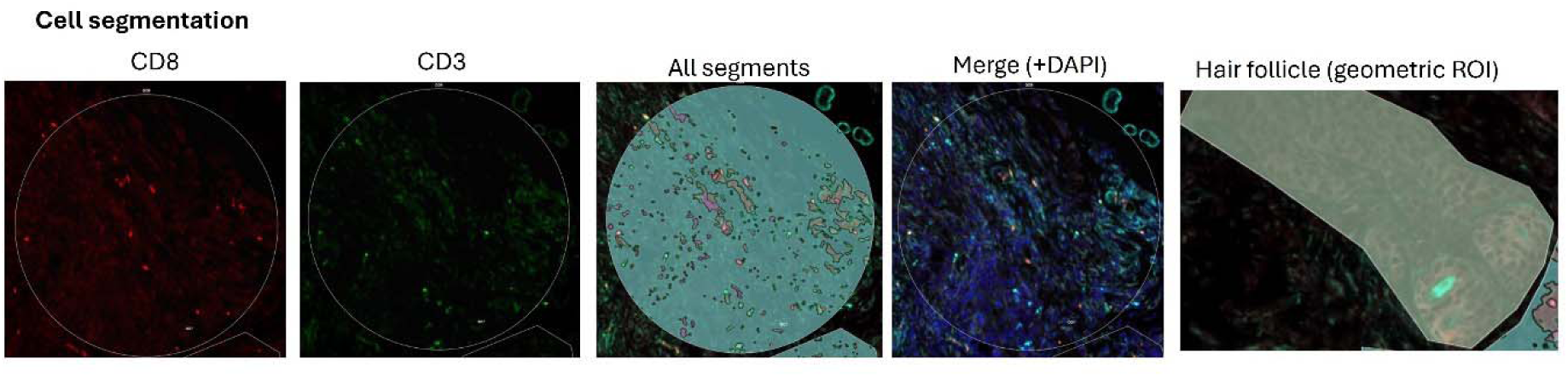
Example cell segmentation used for region of interest selection.

**Figure S4.**
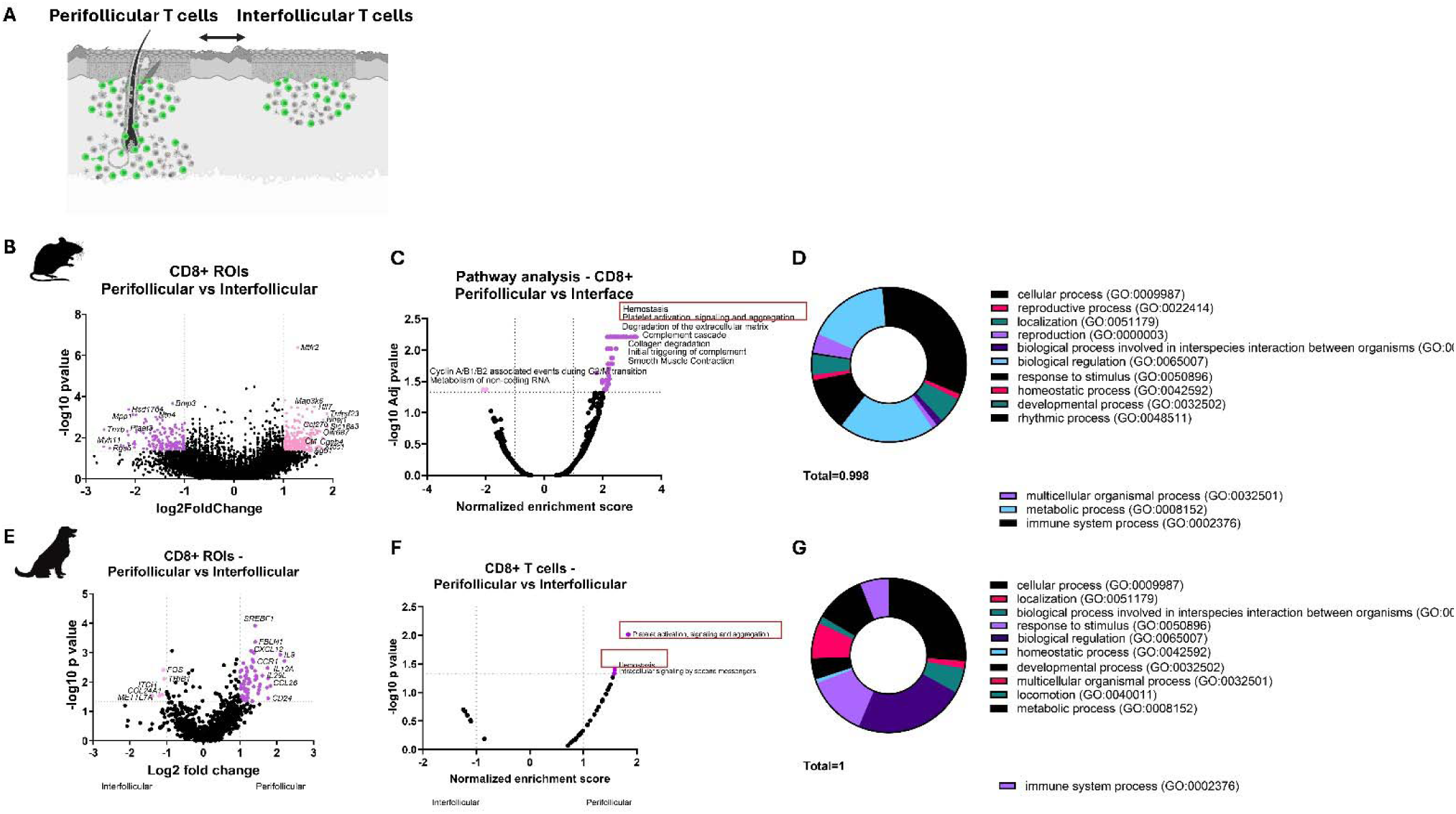
Comparative spatial transcriptomics of mouse and dog perifollicular versus interfollicular CD8+ T cells reveals hemostasis and platelet activation gene ontology terms are conserved in hair follicle associated T cells. A. BioRender schematic of peri versus interfollicular T cell comparison. B. Mouse peri versus interfollicular CD8+ T cell gene expression. C. Pathway analysis of mouse CD8+ T cells reveals hemostasis and platelet activation pathways as the top upregulated terms in perifollicular T cells. D. Biological process GO terms as determined by PantherDB. E. Dog peri versus interfollicular CD8+ T cell gene expression. F. Pathway analysis reveals hemostasis and platelet terms are conserved in perifollicular dog CD8+ T cells. G. GO terms as determined by Panther DB.

**Table S1.**
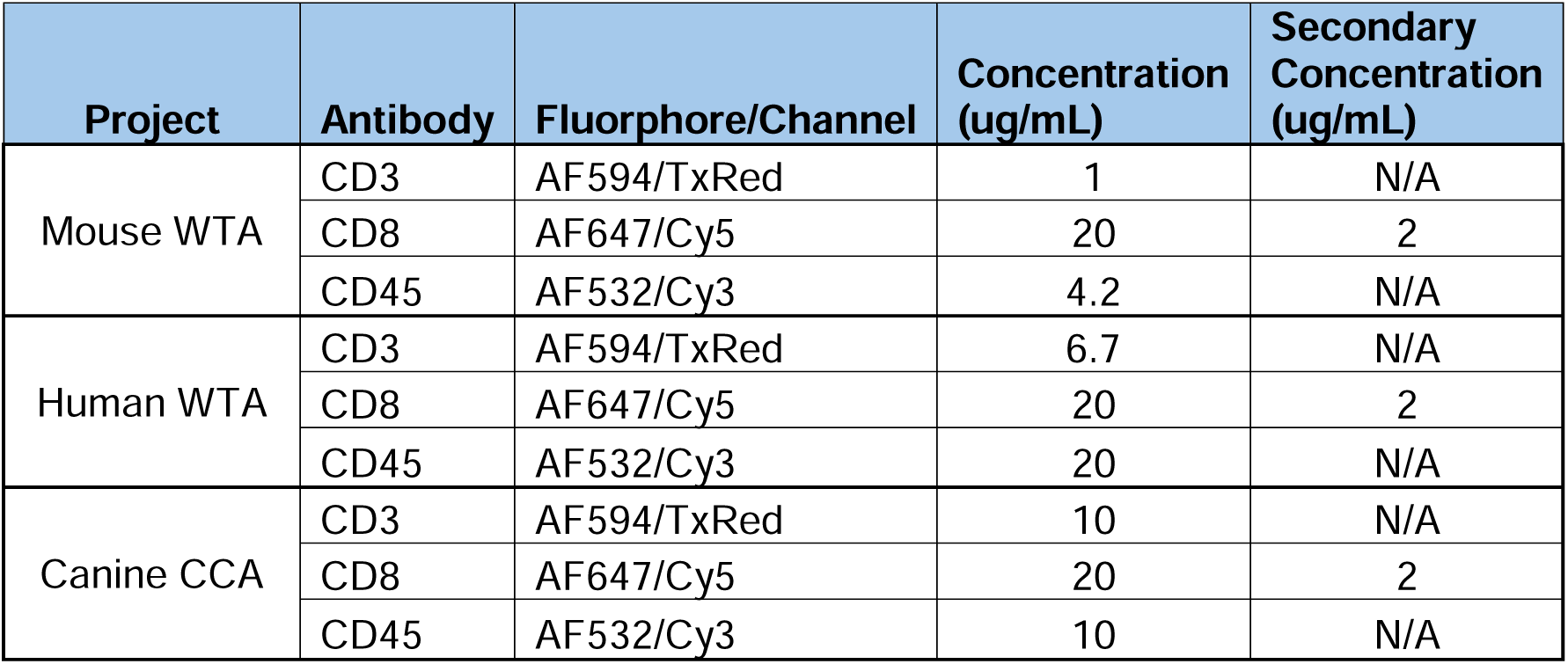
Antibodies used in spatial transcriptomics assays.

| Project | Antibody | Fluorophore/Channel | Concentration (ug/mL) | Secondary Concentration (ug/mL) |
| --- | --- | --- | --- | --- |
| Mouse WTA | CD3 | AF594/TxRed | 1 | N/A |
|  | CD8 | AF647/Cy5 | 20 | 2 |
|  | CD45 | AF532/Cy3 | 4.2 | N/A |
| Human WTA | CD3 | AF594/TxRed | 6.7 | N/A |
|  | CD8 | AF647/Cy5 | 20 | 2 |
|  | CD45 | AF532/Cy3 | 20 | N/A |
| Canine CCA | CD3 | AF594/TxRed | 10 | N/A |
|  | CD8 | AF647/Cy5 | 20 | 2 |
|  | CD45 | AF532/Cy3 | 10 | N/A |

**Table S2.** Antibodies used in mouse flow cytometry panels.

| Hair follicle panel* |  |  |  |  |  |  |
| --- | --- | --- | --- | --- | --- | --- |
| Fluorophore (Color) | marker | clone | catalog # | lot # | RRID | Reactivity |
| **BV570** | CD45 | 30-F11 | 103136 | B419084 | AB_2562612 | Mouse |
| **PE/Cy7** | CD31 | 390 | 102418 | B169004 | AB_830757 | Mouse |
| **PerCP/Cy5.5** | Ep-CAM (CD326) | G8.8 | 118220 | B378718 | AB_2246499 | Mouse |
| **PE/Dazzle 594** | CD200 | OX-90 | 123820 | B319841 | AB_2832446 | Mouse |
| **PE/Dazzle 594** | CD200 | OX-90 | 123819 | B319840 | AB_2832445 | Mouse |
| **APC** | CD200 | OX-90 | 123810 | B203311 | AB_10900447 | Mouse |
| **APC/Fire™ 750** | CD34 | MEC14.7 | 128614 | B312187 | AB_2715985 | Mouse |
| **AF700** | Sca-1 | D7 | 108142 | B377056 | AB_2565959 | Mouse |
| **BV750** | CD3 | 17A2 | 100249 | B394145 | AB_2734148 | Mouse |
| **BV650** | CD11b | M1/70 | 101259 | B376138 | AB_2566568 | Mouse/Human |
| **BV711** | Thy-1.1 | OX-7 | 202539 | B362072 | AB_2562645 | Rat/Mouse |
| **BV711** | CD19 | 6D5 | 115555 | B298801 | AB_2565970 | Mouse |
| **BV421** | F4/80 | BM8 | 123137 | B362959 | AB_2563102 | Mouse |
| **BV785** | F4/80 | BM8 | 123141 | B331869 | AB_2563667 | Mouse |
| **BV785** | CD62L | MEL-14 | 104440 | B272550 | AB_2629685 | Mouse |
| **BV605** | MHC II (I-A/I-E) | M5/114.15.2 | 107641 | B294302 | AB_2565975 | Mouse |
| **PE** | TCR Vβ5.1, 5.2 | MR9-4 | 139504 | B290269 | AB_10613279 | Mouse |
| **PE** | CD200R | OX-110 | 123916 | B345476 | AB_2810379 | Mouse |
| **FITC** | C3aR | 74 | MA5-17475 | YC3877592 | AB_2538865 | Rat |
| **Memory T cell panel** |  |  |  |  |  |  |
| **Fluorophore (Color)** | **marker** | **clone** | **catalog #** | **lot #** | **RRID** | **Reactivity** |
| **BV750** | CD3 | 17A2 | 100249 | B394145 | AB_2734148 | Mouse |
| **BV570** | CD45 | 30-F11 | 103136 | B419084 | AB_2562612 | Mouse |
| **BV421** | CD69 | H1.2F3 | 104545 | B384579 | AB_2686969 | Mouse |
| **BV605** | CD103 | 2 E 7 | 121433 | B387609 | AB_2629724 | Mouse |
| **BV711** | CXCR6 | SA051D1 | 151111 | B352518 | AB_2721558 | Mouse |
| **PE/Cy7** | IL-2Rβ / CD122 | TM-β1 | 123216 | B306704 | AB_2562895 | Mouse |
| **BV650** | IL-7Rα / CD127 | A7R34 | 135043 | B399747 | AB_2629681 | Mouse |
| **PE/Cy7** | CD200R3 | Ba103 | 142211 | B297487 | AB_2814045 | Mouse |
| **APC** | IL-15Rα / CD215 | DNT15R A | 153505 | B309655 | AB_2734220 | Mouse |
| **APC/Fire750** | CCR1 | S15047E | 152512 | B347940 | AB_2566370 | Mouse |
| **BV785** | CD62L | MEL-14 | 104440 | B400007 | AB_2629685 | Mouse |
| **PE** | CD200R | OX-110 | 123908 | B331884 | AB_2561942 | Mouse |
| **Alexa Fluor 700** | CD62L | MEL-14 | 104426 | B497511 | AB_493568 | Mouse |
| **APC/Fire750** | CD44 | IM7 | 103062 | B375202 | AB_2563822 | Mouse |

**\*Different markers were used in different iterations of the panel for BV711, BV785, PE and PE/Dazzle 594.**

**Table S3.** Human CLE biopsies used for spatial transcriptomics.

| Slide & position | Age | Sex | Race/Ethnicity | Site | Diagnosis | Presentation | Serology | Treatment |
| --- | --- | --- | --- | --- | --- | --- | --- | --- |
| A-top | 48 | M | NK | Upper central Back | Discoid Lupus Erythematosus | NK | NK | NK |
| A-middle | 23 | F | African American | Left postauricular | Acute discoid lupus | "Charcoal" hyperpigmented patchy areas over ears | NK | NK |
| A-bottom | 23 | F | African American | Right Upper Arm | Acute discoid lupus | "Charcoal" hyperpigmented patchy areas over ears and skin lesions suspicious of connective tissue disease or vasculitis | NK | NK |
| B-top | 42 | M | NK | Right nasolabial fold | SCLE | Mottled, hyperpigmented, well-demarcated annular patch with superimposed atrophic macules | NK | NK |
| B-middle | 51 | M | African American | Right arm | Cutaneous lupus | Hyperpigmented scaly patches and plaques on face, scalp, legs, arms, trunk | Positive C3, C4, LDH, CK. Negative ANA, anti-dsDNA, lupus anticoagulant | NK |
| B-bottom | 37 | F | African American | Scalp | Discoid Lupus Erythematosus | Erythematous plaque with rim of hyperpigmentation on frontal scalp for 6 weeks | NK | NK |
| C-top | 26 | F | African American | Left arm | SLE | Photosensitive eruptions | NK | NK |
| C-middle | 47 | M | African American | Chest | Tumid form of Lupus | NK | anti-dsDNA negative | Hydroxychloroquine, Mycophenolate |
| C-bottom | 47 | M | African American | Left arm | Tumid form of Lupus | NK | anti-dsDNA negative | Hydroxychloroquine, Mycophenolate |
| D-top | 23 | F | African American | Upper Back | Discoid Lupus Erythematosus | Sun-exposed plaque | anti-dsDNA negative, anti-RNP positive, anti-smith positive, | Hydroxychloroquine |
| D-middle | 33 | F | African American | Left Upper Back | Lupus tumidus | No PMH presents with urticarial papules on upper extremity | NK | NK |
| D-bottom | 55 | F | NK | Right Upper Back | Drug-induced lupus | Erythematous macules/patches and hyperpigmentation on B/L forearms, back, forehead, chest. | NK | NK |

\*NK = not known

## Acknowledgments

We thank Ann Rothstein for TLR9KO mice and Michael Rosenblum for K5 TGO mice. We thank Frane Banovic for insightful discussions of canine DLE, Nazgol-Sadat Haddadi, Khashayar Afshari and Danny Kwong for technical assistance, Jayme Heywosz and Tammy Hayes for DSP sectioning, and Maria Zapp and Ellie Kittler for sequencing. We also thank William Petrides for designing and 3D printing the mouse ear stage, and Clement David for GeoMX software troubleshooting.

## Data availability

All data are presented in the manuscript and supplement.

Mouse microarray data are deposited on Gene Expression Omnibus under accession # GSE285892.

Canine bulk FFPE RNAseq data are deposited on Sequence Read Archive under accession “PRJNA1119020 : RNA sequencing of canine discoid lupus erythematosus and healthy margin skin biopsies”.

Spatial transcriptomics datasets are deposited on Sequence Read Archive under accessions “PRJNA1156903 : Spatial transcriptomics of Mouse Cutaneous Lupus Erythematosus and Primary Cicatricial Alopecia Skin”, and “PRJNA1156903 : Spatial transcriptomics of Canine Cutaneous Lupus Erythematosus Skin”.

Primary data from humans will be made available upon reasonable request under Material Transfer Agreements.

## CRediT statement

Conceptualization – JMR

Methodology - CB, MA

Software - N/A

Validation - RA, WA, LY, ES, HB

Formal analysis - UYA, AR, JMR

Investigation - UYA, TC, SS, MD, QT, MO, FB,

JMR Resources - BS, ASB, CF, CB, AK, RMA, JMR

Data Curation - UYA, JMR

Writing - Original Draft - UYA, JMR

Writing - Review & Editing - all authors Visualization - UYA, JMR

Supervision - JMR, CF, CB

Project administration - JMR

Funding acquisition - JMR, CB, WA

