## Supplementary figures and images for "Comparative spatial transcriptomics of hair follicle-T cell interactions in mouse, dog and human reveals conserved drivers of primary cicatricial alopecia"

### Fig S1

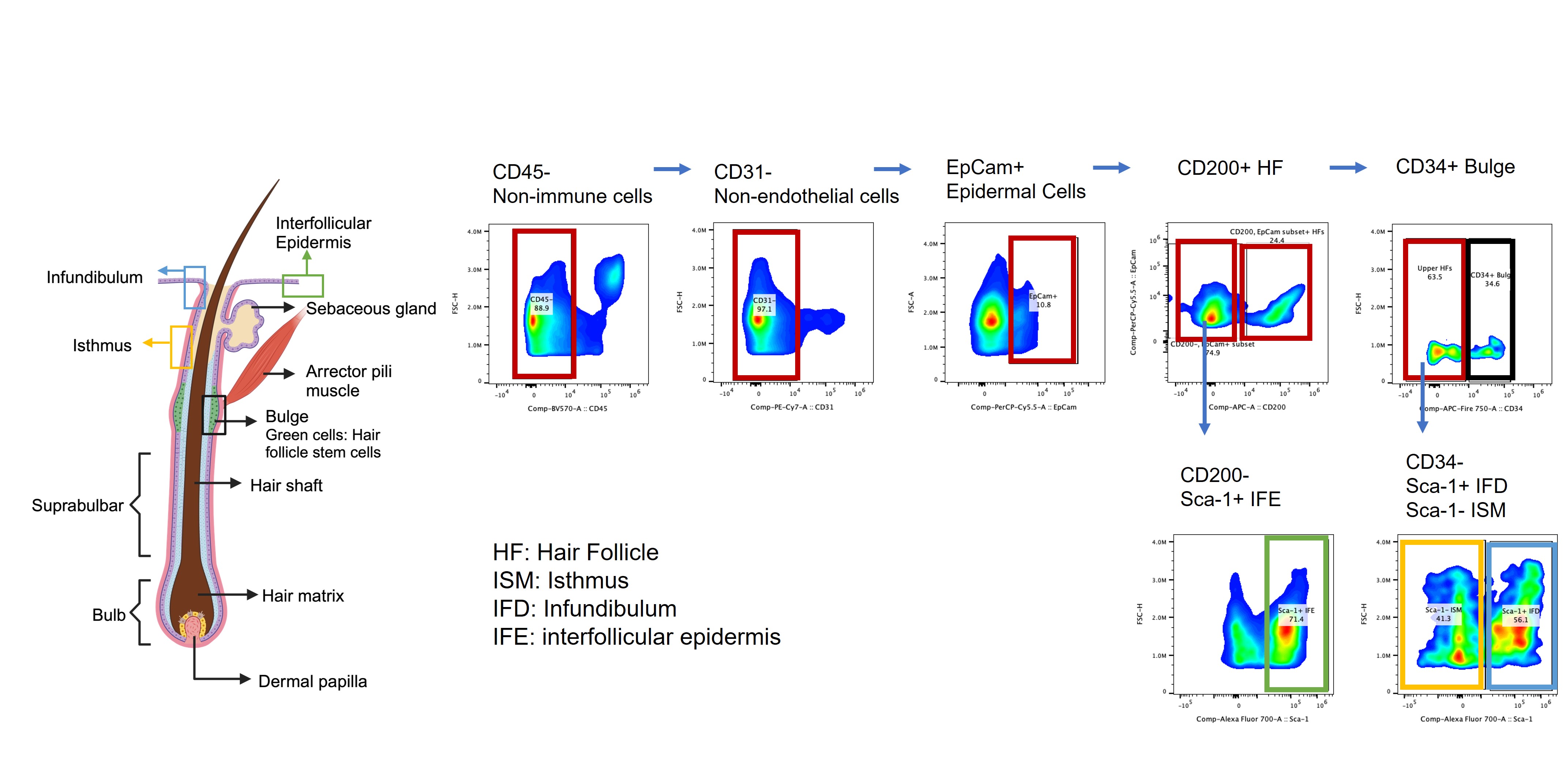

### Fig S2

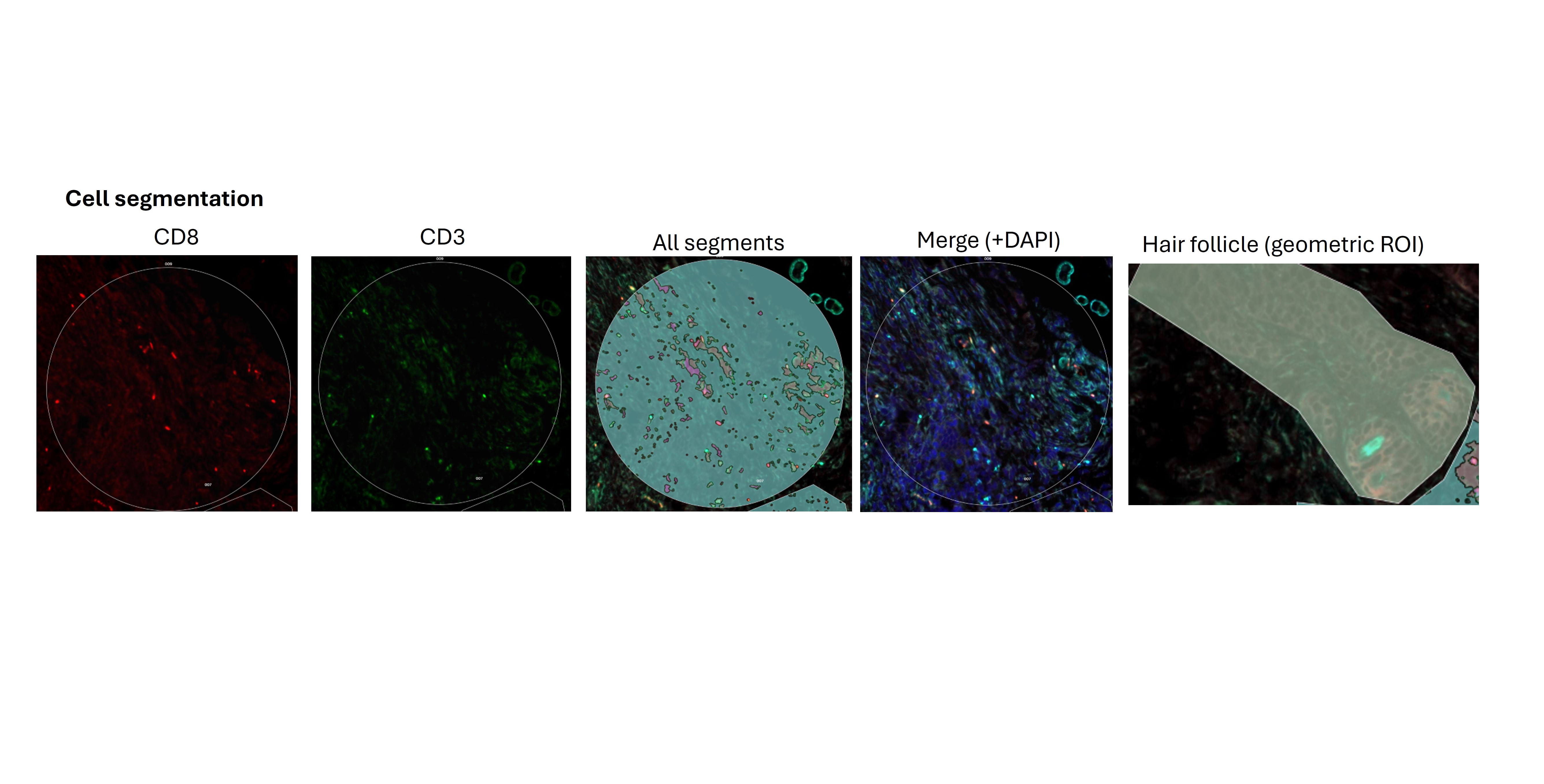

### Fig S3

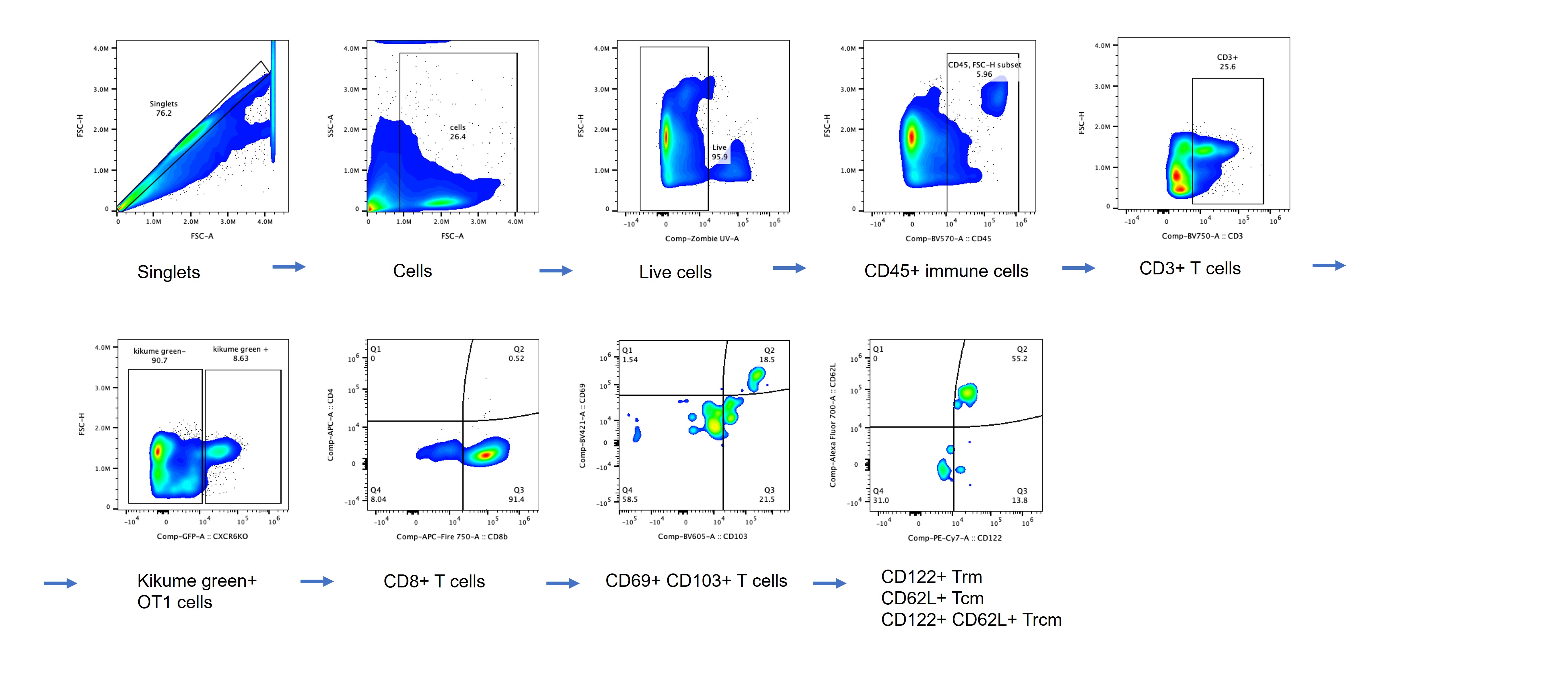

### FIg S4

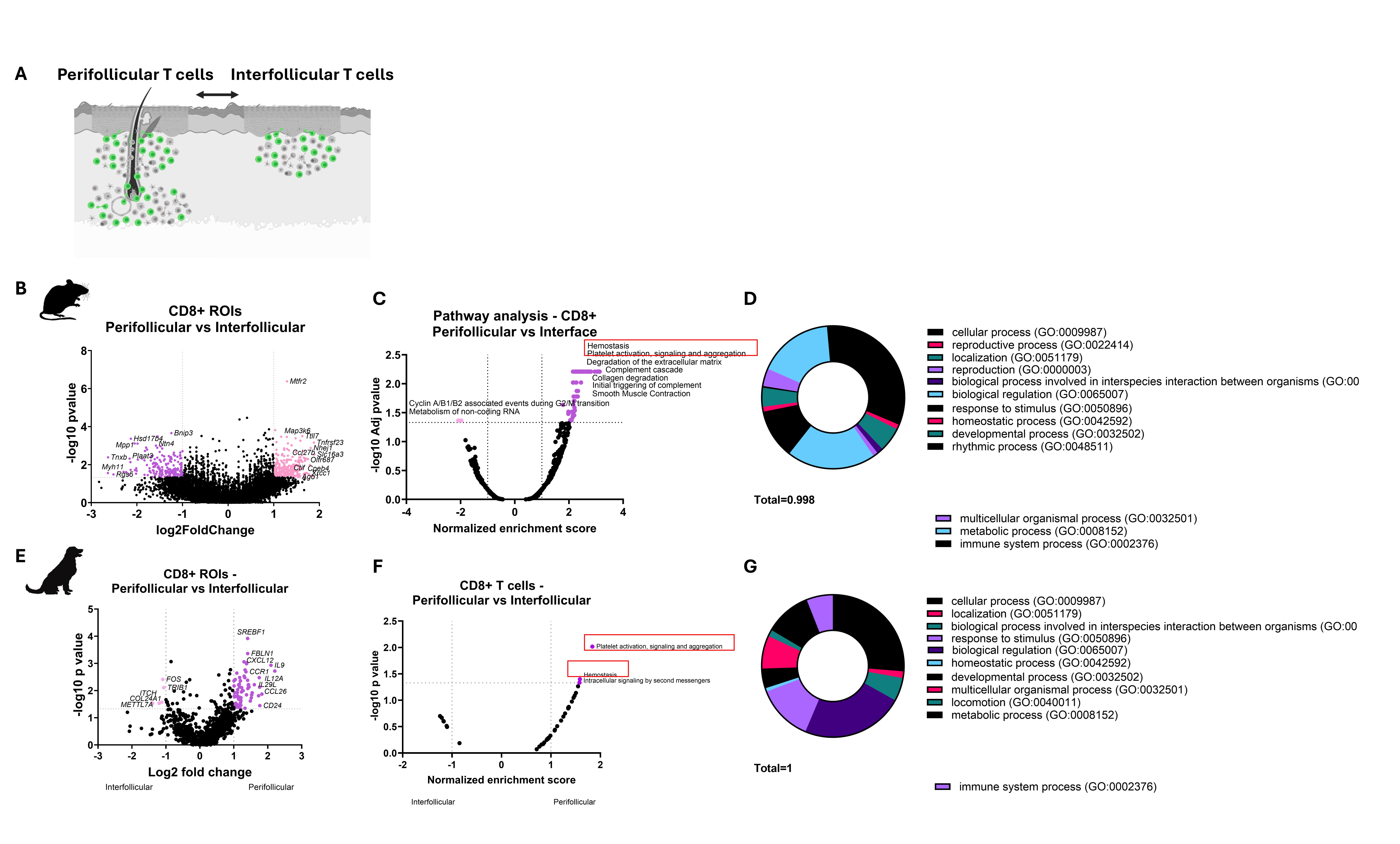
